# A new approach to integrate phylogenetic structure and partner availability to study biotic specialization in ecological networks

**DOI:** 10.1101/2021.08.04.454912

**Authors:** Carlos J. Pardo-De la Hoz, Ian D. Medeiros, Jean P. Gibert, Pierre-Luc Chagnon, Nicolas Magain, Jolanta Miadlikowska, François Lutzoni

**Affiliations:** Department of Biology, Duke University, Durham, North Carolina, 27708, United States of America; Département des Sciences Biologiques, Université de Montréal, Montréal, Québec, H1X 2B2, Canada; Biologie de l’évolution et de la conservation, Université de Liège, Liège, 4000, Belgium

**Keywords:** Community phylogenetics, coral symbiosis, host-parasite interaction, lichen symbiosis, mutualism, seed dispersal

## Abstract

Biotic specialization holds information about the assembly, evolution and stability of biological communities. Phylogenetic diversity metrics have been used to quantify biotic specialization, but their current implementations do not adequately account for the availability of the interacting partners. Also, the overdispersed pattern of phylogenetic specialization has been misinterpreted as an attribute of generalists. We developed an approach that resolves these issues by accounting for partner availability to quantify the phylogenetic structure of specialization (i.e., clustered, overdispersed, or random) in ecological networks. We showed that our approach avoids biases of previous methods. We also implemented it on empirical networks of host–parasite, avian seed-dispersal, lichenized fungi– cyanobacteria and coral–dinoflagellate interactions. We found a large proportion of taxa that interact with phylogenetically random partners, in some cases to a larger extent than detected with an existing method that does not account for partner availability. We also found many taxa that interact with phylogenetically clustered partners, while taxa with overdispersed partners were rare. Our results highlight the important role of randomness in shaping interaction networks, even in highly intimate symbioses, and provide a much-needed quantitative framework to assess the role that evolutionary history and symbiotic specialization play in shaping patterns of biodiversity.

## Introduction

Species interactions display multiple levels of biotic specialization that impact the evolution, assembly and stability of biological communities (Guimarães *et al*., 2011; Poisot *et al*., 2011; Chomicki *et al*., 2019). Ecologists typically characterize these levels of biotic specialization by quantifying two aspects of a species’ niche: the niche breadth (e.g., the number of species with which they interact; partner breadth henceforth), and the intensity of the interactions with those partners (Colwell & Futuyma, 1971; Hurlbert, 1978; Futuyma & Moreno, 1988; Pinheiro *et al*., 2016). These interactions are often studied as networks, with nodes representing species and links representing ecological interactions (Jordano, 1987; Guimarães *et al*., 2006). The structural features of these networks contain information on at least one of the two aspects of biotic specialization (Blüthgen *et al*., 2008). For example, the total number of partners of a species in a network, or node degree, characterizes partner breadth (Jordano *et al*., 2002), while the distribution of interaction frequencies across partners (interaction strength sensu Vázquez *et al*., 2007) provides information on the intensity of partner use. Additionally, network structural features such as the presence of subsets of species that interact preferentially with one another, also known as modularity, provide information about the level of partner sharing between species in a community (Olesen *et al*., 2007). Thus, interaction networks are a powerful framework to study the spatiotemporal factors shaping biotic specialization (Blüthgen *et al*., 2007; Schleuning *et al*., 2012; Fortuna *et al*., 2020).

One limitation for characterizing partner breadth in interaction networks is the use of discrete counts for the number of partners of a species. Even if two species associate with the same number of partners, one of them might be specialized on partners with a narrower range of traits (Dehling *et al*., 2020). Furthermore, most symbioses involve microbes for which species boundaries are unclear, making it difficult to quantify the number of partners (Toju *et al*., 2014; Magain *et al*., 2017a; Põlme *et al*., 2018). One alternative is to measure the range of partner traits directly. For instance, Junker *et al*. (2013) measured the range of flower traits in plants visited by arthropod species as a proxy for the partner breadth of the arthropods in a pollination network. However, many interactions are mediated by traits that are unknown or difficult to measure. In these cases, phylogenetic relatedness is a useful alternative for characterizing the partner breadth of a species (Faith, 1992; Webb, 2000) because traits tend to be phylogenetically conserved (reviewed in Swenson, 2013). For example, phylogenetic diversity metrics are often used to quantify the specialization of parasites on their hosts (Cooper *et al*., 2012; Lane *et al*., 2014; Esser *et al*., 2016; Doña *et al*., 2018). Under this approach, parasites are considered more specialized if they associate with hosts that are more closely related than expected by a null model (Poulin *et al*., 2011).

Phylogenetic diversity metrics used to quantify biotic specialization can be modified to integrate information about the intensity of partner associations (Webb, 2000; Webb *et al*., 2008; Miller *et al*., 2017). This is done by using the interaction frequencies between the partners of the focal species as weights for the phylogenetic distances between the partners (Webb *et al*., 2008). One caveat of this approach is that changes in partner availability can change interaction frequencies and alter the number of partners for a focal species when interactions occur randomly (Fig. **1a–c**; Lessard *et al*., 2012; Poisot *et al*., 2015). As a result, a species may appear specialized on a set of closely related partners (i.e., phylogenetically clustered) only because the most available partners happened to be closely related (Fig. **1b**). Previous simulation studies have revealed multiple scenarios where failure to account for the distribution of species abundances (analogous to partner availability) results in biased estimates of phylogenetic diversity (Kembel, 2009; Miller *et al*., 2017). One way to account for this problem is to compare observed values of phylogenetic diversity to a null distribution generated by drawing partners from a pool of species in proportion to their availability, instead of drawing them with equal probabilities (Kembel, 2009; Jorge *et al*., 2014). However, this null model is also biased (Fig. **S1**), and results in an overestimation of non-random phylogenetic structure with increasing interaction frequencies, as evidenced by Jorge *et al*. (2017) for networks of plant–herbivore interactions.

**Figure 1.**
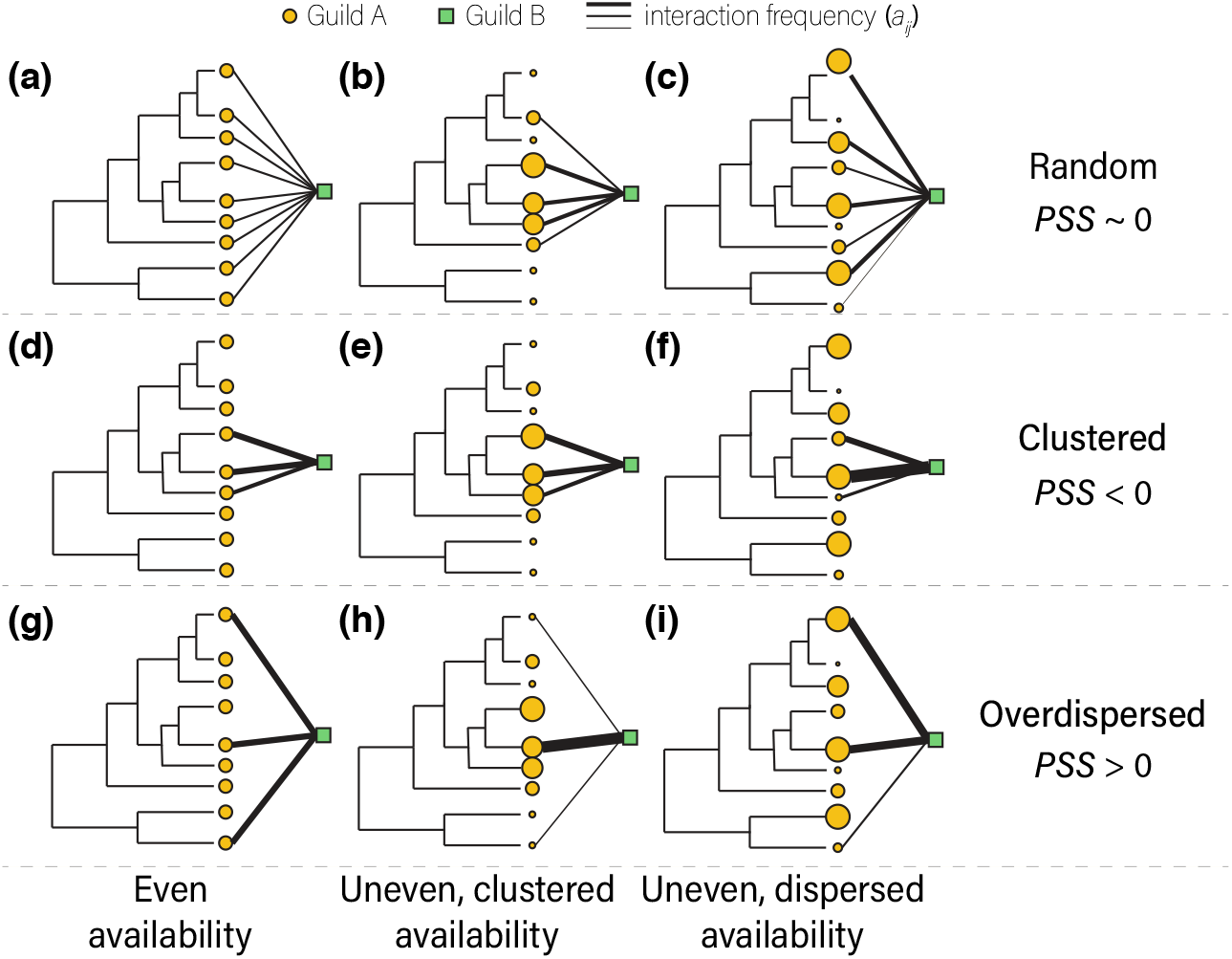
Schematic representation of three possible patterns of phylogenetic structure and their corresponding Phylogenetic Structure of Specialization (PSS) values (random [**a–c]**, clustered [**d–f**], and overdispersed [**g–i**]) across three different patterns of partner availability from guild A that interact with one species from guild B. The size of the orange circles represents the relative availability of each member of guild A in nature, or frequencies in an interaction matrix as a proxy for availability in nature.

The statistical biases of current approaches limit our ability to detect the patterns of biotic specialization in ecological networks and the processes that shape them. To solve this issue, we developed an approach to infer the phylogenetic structure of specialization in bipartite ecological networks while accounting for species availability. We designed our approach to overcome the biases of existing methodologies. We conducted simulations to detect potential biases of this new approach, and illustrate its use with four bipartite networks from the literature for which molecular phylogenetic trees were available for both sets of partners. We propose a conceptual framework to interpret phylogenetic structural patterns of biotic specialization in ecological networks (Figs **1, 2**) that enables the exploration of putative ecological and evolutionary processes generating these patterns.

## Materials and Methods

### 1. The phylogenetic structure of specialization (PSS) index

Phylogenetic diversity indices used to measure specialization are Standardized Effect Sizes (SES; Poulin *et al*., 2011; Miller *et al*., 2017). As such, these indices compare observed values of a phylogenetic diversity metric to a null distribution (i.e., SES = (null_mean_ – observed)/null_sd_). A strategy to account for availability is to generate the null distribution by calculating the phylogenetic diversity of sets of partners that are drawn from the pool of partner species in proportion to their availability (Jorge *et al*., 2014). However, this procedure often yields null sets with a different number of partner species compared to the observed number of partners of the focal species. If the null and observed values of phylogenetic diversity are not based on the same number of species, the SES will be biased (Jorge *et al*., 2017). This is because phylogenetic diversity metrics are not independent from the number of species upon which they are calculated (Fig. **S1**; Appendix **S1**). We avoid this problem by accounting for partner availability in the calculation of the phylogenetic diversity metric itself. This ensures that the SES is calculated with observed and null values that are based on the same number of species.

Specifically, we use Kullback-Leibler distances (Kullback & Leibler, 1951) to determine the deviations of observed interaction frequencies from a null expectation where interaction frequencies are only driven by the availability of each partner (Blüthgen *et al*., 2006). We incorporate these deviations as weights into the calculation of the mean pairwise phylogenetic distance, a common metric of phylogenetic α-diversity (*wMPD*; Webb *et al*., 2008). Because the availability of interacting partners has already been accounted for in the observed values of the raw metric of phylogenetic diversity, we can generate a null distribution of phylogenetic distances by shuffling taxa at the tips of the partner phylogeny. Then, observed values are compared to null values that are simulated with the same distribution of interaction frequencies and total number of partners. We provide details of the calculation of the PSS index below.

First, we show the calculation of the Kullback-Leibler distances that are used to account for the availability of the partners. We let **I** be an interaction matrix with *r* species in the rows (guild A), and *c* species in the columns (guild B). Each element *a*_*ij*_ of **I** represents the interaction frequency between species *i* and species *j*, such that:

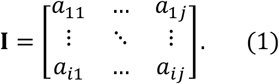

Let A_*i*_ be the sum of interaction frequencies recorded for species *i*,

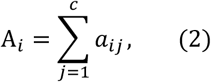

and *m* the total number of interactions in the matrix (i.e., the sum of interaction frequencies across both rows and columns),

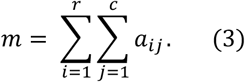

Let *q*_*j*_ be the proportion of the sum of interaction frequencies of species *j* to the total number of interactions in the matrix,

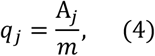

and the proportion of the interaction frequency between species *i* and *j*, relative to the sum of interaction frequencies of species *i*, P’_*ij*_, be written as:

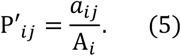

Then, the species level specialization,*d*_*i*_,, can be defined as the sum of the Kullback-Leibler distances between P’_*ij*_ and *q*_*j*_ (Blüthgen *et al*., 2006), defined as:

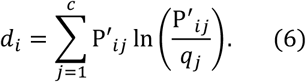

These distances tell us how much the interaction frequencies of species *i* deviate from a null model where all partner species are used in proportion to their frequency in the entire interaction matrix.

To calculate the phylogenetic diversity metric, we defined M_*i*_ as the set of species that associate with species *i*, and **D**_*i*_ as a symmetric matrix of pairwise phylogenetic distances between all species that belong to the set M_*i*_:

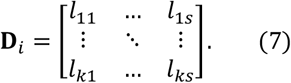

The weighted mean pairwise phylogenetic distance (Webb *et al*., 2008) of the species that associate with *i, wMPD*_*i*_, is defined as:

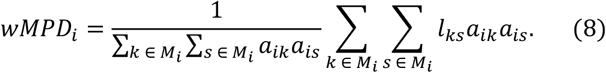

This is a mean of the phylogenetic distances between a set of species. It takes larger values for sets of distantly related species and smaller values for sets of closely related species. This mean is weighted by the interaction frequencies of species *i, a*_*ik*_*a*_*is*_, and is used to compute the Net Relatedness Index (NRI) as an SES, which is commonly used to quantify phylogenetic specialization. This metric can yield biased estimates of the phylogenetic structure, especially when the availability of the partners has phylogenetic signal (Kembel, 2009; Miller *et al*., 2017). To remove this effect, for a species *k* that associates with *i* and belongs to the set M_*i*_, we defined a KL_*k*_ factor:

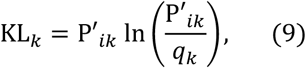

which corresponds to an element of the sum in equation 6. When interactions are random, the proportion of interactions of *i* with *k*, P’_*ik*_, should converge to the availability of *k*, or *q*_*k*_. Therefore, the ratio between these two parameters tends toward 1, and KL_*k*_ will approach 0. Conversely, when interactions are non-random, the proportion of interactions of *i* with *k*, P’_*ik*_, is larger than the availability of *k, q*_*k*_, and KL_*k*_ becomes larger than 0. In equation 9, the availability parameter, *q*_*k*_, can be inferred from the interaction matrix as shown in equation 4 (as a proxy for partner availability), or it can be determined empirically.

The latter is preferable when data are available (Jorge et al., 2014).

We replaced the interaction frequencies in equation 8, *a*_*ik*_*a*_*is*_, with the KL factors from equation 9 to compute a version of the *MPD* for species *i, klMPD*_*i*_, that is weighted by the KL factors, instead of the interaction frequencies:

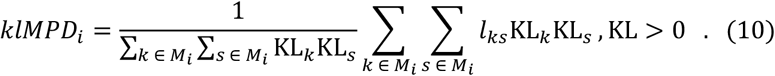

This mean of pairwise distances is now corrected for the availability of the partners through the KL weights. It takes larger values for species that interact non-randomly with sets of more distantly related species, and it is undefined (0 in both numerator and denominator) for a species that interacts with its partners at the exact frequency that those partners are available (i.e., P’_*ik*_ = *q*_*k*_ for every *k* that belongs to M_*i*_). However, the scenario where the index is undefined is extremely unlikely in natural networks and was never found in simulated networks. Equation 10 may result in negative values when KL factors are ≥ 0. Consequently, we only consider partners for which the KL factors are > 0. This means that we only use phylogenetic distances among partners that seem to be preferentially selected by the focal species. See Appendix **S2** for a discussion on why this does not affect the behavior of the index.

For each species *i*, PSS_*i*_ is calculated as the difference between the observed value of *klMPD*_*i*_ and the mean 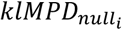 values of a null distribution—obtained by randomizing the tips of the phylogenetic tree of the partner species, which maintains the observed total number of partners—divided by the standard deviation of such null values. PSS is thus an SES, with values close to 0 indicating that the partners of a focal species lack phylogenetic structure, and negative or positive values indicating phylogenetic clustering or overdispersion, respectively (Fig. **1**). The PSS index is thus:

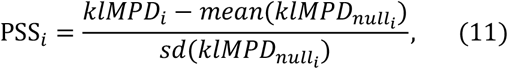

where the null distribution is generated from 999 iterations of random taxon shuffling using the empirical phylogeny. As a guild-level measure of the PSS, we took the mean of the SES from all species in the rows (guild A) weighted by the interaction frequencies of each species:

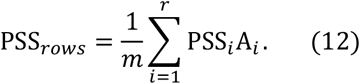

PSS scores are independent from Blüthgen’s *d’* (equation 6). For example, a link between two species with high *d’* values would occupy the specialist-specialist region in Fig. **2a** (they do not need to be reciprocally specialized) based on their interaction frequencies. However, PSS (Fig. **2c**) integrates both partner availability, as in *d’* (Fig. **2a**), and phylogenetic structure, as in NRI (Fig. **2b**). Therefore, the same interacting pair can occupy any region in the PSS–PSS space depending on the phylogenetic relatedness of their preferred partners (Fig. **2c**). With *d’*, the terms ‘specialist’ and ‘generalist’ refer exclusively to the interaction frequencies relative to the availability of the partners (Blüthgen *et al*., 2006). For example, a species can have *d’* = 0 (generalist) regardless of the number of partners with which it associates, as long as that species interacts with those partners in the same proportion as they are available.

**Figure 2.**
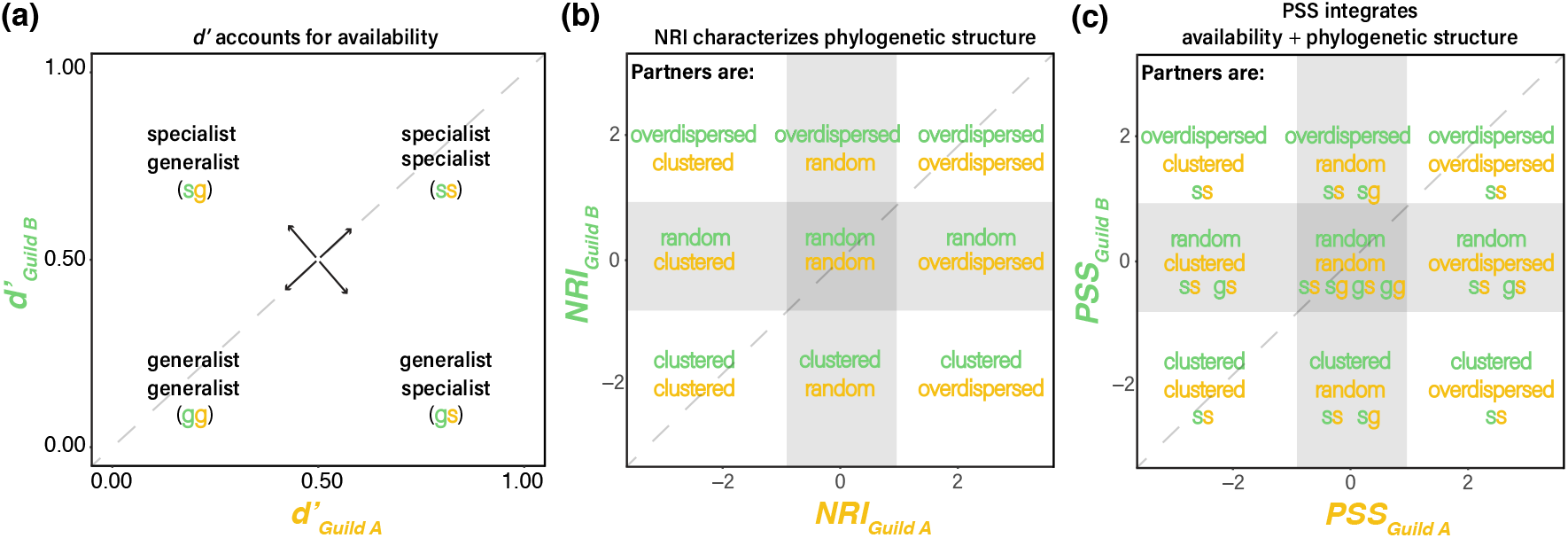
Schematic representation of the variation of *d’* (**a**), NRI (**b**), and PSS (**c**) in a co-specialization profile of a bipartite interaction network involving guilds A and B. The X axis represents *d’*, NRI, or PSS values for the species in the rows of an interaction matrix (guild A), and the y axis represents the *d’*, NRI, or PSS values for the species in the columns of an interaction matrix (guild B). The diagonal dashed lines represent specialization symmetry. The shaded gray areas in panels **b** and **c** show the NRI and PSS space where clustering or overdispersion are not significantly different from a random pattern of phylogenetic structure (Fig. **1a–c**). These thresholds flanking the shaded gray areas (−1,1) represent the 95% confidence interval of the null distribution of NRI or PSS simulated under each dataset. Each region of the NRI–NRI and PSS–PSS space is labeled with the phylogenetic structure of the partner guilds. The color of the labels relates to guild A or B in accordance with Fig. **1**. Note that NRI does not account for availability, and therefore specialization as defined for *d’* cannot be mapped to the NRI–NRI space. In contrast, PSS integrates both availability and phylogenetic structure. For example, a species from guild B with PSS = -2, is considered a specialist that associates with phylogenetically clustered partners from guild A (see also Fig. **1d–f**).

### 2. Why use the mean pairwise phylogenetic distance (MPD) for PSS?

Metrics of phylogenetic α-diversity fall into one of three groups based on what they quantify (Swenson, 2014; Miller *et al*., 2017; but see Tucker *et al*., 2017 for an alternative classification): i) the mean relatedness among species, such as the MPD (Webb, 2000); ii) relatedness of species to their closest relatives, such as the Mean Nearest Taxon Distance (Webb, 2000); or iii) distances to a centroid, such as Faith’s Phylogenetic Distance (PD; Faith, 1992). Our approach to account for availability could be coupled with any metric of phylogenetic α-diversity, i.e., by incorporating the KL factors as weights in the calculations of the mean. However, there are caveats associated with specific types of metrics. For example, metrics from group ii only provide insights about fine-scale phylogenetic structure because only the closest relatives are considered. Additionally, metrics from group iii do not provide information about phylogenetic structure because taxa may have similar distances to a phylogenetic centroid, regardless of how related they are to each other. The use of MPD for our PSS index allows the detection of three different phylogenetic structural patterns of specialization (random, clustered and overdispersed; Fig. **1**).

However, MPD can only be calculated if one lineage is interacting with at least two partners. This is problematic in highly specialized symbiotic systems where many species interact with one partner. Even if we assume that a species with one partner has a MPD of 0, as in previous studies (Jorge *et al*., 2014, 2017), PSS cannot be calculated because the null distribution would also be estimated with one partner, resulting in an undefined PSS with the numerator and denominator of equation 11 being equal to 0. We solved this problem by assuming the existence of a sister taxon (by adding one terminal branch in the phylogeny) to the recorded single partner of the focal species. This new sister taxon is joined to the original partner with a branch length that is half the minimum pairwise distance recorded between any pair of species in the phylogeny of the partners. The observed interaction frequencies of the original partner are divided by two, and we assume the new sister partner also has the interaction frequencies of the original partner divided by two. This keeps the relationship between interaction frequencies and availability (i.e., P’_*ik*_/*q*_*k*_ in equation 9) constant, thus retaining the appropriate proportions of the KL factors. This new sister taxon is only added for the calculation of K@CDE_#_ (equation 10) for species with one partner and does not affect species that have more than one partner.

### 3. Testing the PSS index using simulations

#### 3.1 Varying dimensions and total number of interactions

We simulated random matrices using the genweb() function in the R package *bipartite* (Dormann *et al*., 2008), which relies on three parameters: number of columns (N1), number of rows (N2), and average interaction frequency per link (dens). These values are used to calculate the sum of all interaction frequencies of the simulated matrix (*m*), as *m* = N1×N2×dens. Then, the simulated matrix of dimensions N1×N2 is populated with *m* interactions, such that the marginal sums of the interaction frequencies in the rows and the columns follow a lognormal distribution. This procedure results in heterogeneous distribution of both the number of links per species and the interaction frequencies per link.

We simulated three sets of random matrices. In the first set, we assessed the values of PSS on random symmetric matrices with dimensions varying from 5×5 to 200×200. The second set consisted of random matrices with unequal numbers of columns and rows varying from 2×20 to 200×20. The average interaction frequency per link (dens) was kept at 2 for all matrix configurations in the first two sets. The third set contained random matrices with increasing total number of interactions (*m =* 5–4000) and fixed dimensions (50×50). We accomplished this by varying “dens” within the genweb() function in *bipartite*. The total number of interactions (*m*) sets a limit to the number of binary links in the simulated matrices because the marginal totals are constrained to follow a lognormal distribution. Therefore, the random matrices in the third set also span a range of connectance values, with matrices with higher *m* having higher binary connectance (Fig. **S2**). Each simulation step consisted of (i) a random matrix simulated as described above, and (ii) a random ultrametric tree, with the matrix columns as taxa generated with the function rcoal() in the R package *ape* (Paradis *et al*., 2004). We then calculated PSS_*rows*_ for each matrix. Each simulation step was replicated 10 times. In addition, each set of simulations was performed three times, with branch length distributions of the random trees drawn from either a lognormal, normal, or uniform distribution. We estimated the rate of Type I error as the proportion of simulated matrices for each parameter value for which the observed *klMPD* value was significantly different (α = 0.05) from the null distribution generated as described for equation 11.

#### 3.2 Varying nestedness and modularity

To determine whether network structural patterns constrain the possible values of PSS, we tested for correlation between PSS on simulated matrices with varying degrees of nestedness and modularity. In nested networks, specialist species tend to interact with a subset of the partners associated with generalist species (e.g., Guimarães *et al*., 2006). In modular networks, groups of species share a set of preferred partners, resulting in compartmentalized networks (e.g., Chagnon *et al*., 2018). In order to simulate matrices with a gradient of nestedness and modularity values, we started by creating a perfectly nested and a perfectly modular 50×50 binary matrix. Then, each simulation step swapped the positions of a 0 and 1 in the matrices, thus adding noise and decreasing nestedness and modularity (Chagnon, 2015). Although we used binary matrices, we treated them as quantitative so that the network structure could be manipulated in a predictable way. After each step of the nestedness simulation, we calculated nestedness using wNODF (weighted nestedness metric based on overlap and decreasing fill) developed by Almeida-Neto *et al*. (2008) and implemented in *bipartite*. After each step of the modularity simulation, we calculated the modularity (Q) using the simulated annealing algorithm developed by Dormann and Strauss (2014) implemented in *bipartite*. For each simulation, we generated a random ultrametric tree with the matrix columns as taxa using the function rcoal() in *ape*. We then calculated PSS_*rows*_ for the matrices of each simulation step. The availability of the partner species (*q*_*j*_) is calculated as if the network were quantitative (see equations 2–4). Even if all interaction frequencies are 0 or 1, our simulation strategy still generates heterogeneity in the availability of partners (*q*_*j*_) and the KL weights (equation 9) that are used to calculate PSS. Each simulation step was replicated 20 times for both modularity and nestedness analyses. As above, each set of simulations was performed three times, with branch length distributions of the random trees drawn from either a lognormal, normal, or uniform distribution. Type I error rates were estimated as described above.

#### 3.3 Can PSS detect clustering and overdispersion?

Because our approach is an index of phylogenetic diversity, it can be used to describe communities without interactions. Therefore, to determine whether PSS can detect phylogenetic structure patterns (Fig. **1**) when they are present (Type II error rate), we used a simulation framework from a recent study that explored the statistical behavior of a comprehensive set of phylogenetic diversity metrics applied to communities (Miller *et al*., 2017). This allowed us to compare PSS to existing methods. This framework simulates arenas where individuals are spatially distributed according to their phylogenetic relatedness (Appendix **S3**). We then sampled the species composition of plots within this arena to create Community Data Matrices (CDM). A CDM is analogous to an interaction matrix, where rows correspond to plots in the arena and columns correspond to species that can be found in the plots. The PSS index was calculated for the plots within these CDMs. Rates of Type II error were calculated at the CDM and plot level as described in Appendix **S3**.

### 4. Empirical networks with phylogenies

We used four bipartite networks from the literature for which molecular phylogenetic trees were available for both sets of partners: (I) mammals-fleas, an antagonistic network of interactions between small mammals and their ectoparasitic fleas (order Siphonoptera) that were sampled in four regions of Slovakia (Javorie mountains, Krupinská Planina plain, Volovské Vrchy mountains, and East Slovakian Lowland; Stanko *et al*., 2002); (II) avian seed-dispersal, a mutualistic network of bird-seed dispersal interactions compiled from studies conducted across multiple localities in the Brazilian Atlantic Forest (Bello *et al*., 2017); (III) cyanolichens, a mutualistic network of interactions between the fungal lichen-forming genus *Peltigera* and their cyanobacterial partners from the genus *Nostoc*, which were recorded at a global scale as part of phylogenetic studies on *Peltigera* (O’Brien *et al*., 2005, 2013; Miadlikowska *et al*., 2014, 2018; Magain *et al*., 2017a,b, 2018; Lu *et al*., 2018; Pardo-De la Hoz *et al*., 2018) and compiled by Chagnon *et al*. (2019); and (IV) corals-dinoflagellates, a mutualistic network of interactions between reef-building corals (order Scleractinia) and their dinoflagellate symbionts (family Symbiodiniaceae) that were recorded as part of multiple community studies of reef-building corals in the Caribbean (LaJeunesse, 2001, 2002, 2005; Garren *et al*., 2006; Thornhill *et al*., 2006, 2009; Kemp *et al*., 2008; Camargo *et al*., 2009; Correa *et al*., 2009; LaJeunesse *et al*., 2009; DeSalvo *et al*., 2010; Finney *et al*., 2010; Green *et al*., 2010) and compiled by Fabina *et al*. (2012) from the GeoSymbio database (Franklin *et al*., 2012). Table **1** shows a summary of these four datasets.

#### 4.1 Computing PSS values

We developed an R package (https://github.com/cjpardodelahoz/pss) with functions to compute PSS using interaction matrices and phylogenetic trees from the interacting species as input. Our script has function dependencies from the R packages *ape, bipartite, picante* and *vegan* (Paradis *et al*., 2004; Dormann *et al*., 2008; Kembel *et al*., 2010; Oksanen *et al*., 2019), and includes code modified from Swenson (2014). For each species in each of these datasets, we calculated node degree, the interaction frequency-based specialization index *d’* (Blüthgen *et al*., 2006), the phylogenetic diversity index NRI (Webb *et al*., 2002), and PSS. The availability parameter (*q*_*k*_; equation 9) was estimated from the interaction frequencies in the matrices for all datasets as indicated in equation 4. We also estimated PSS values for the fleas using the empirical availabilities that were available for the mammals in this dataset (Table **1**; Stanko *et al*., 2002).

## Results

### PSS is independent from basic network features

We calculated PSS across simulated bipartite matrices lacking phylogenetic structure but varying in size, number of rows and columns, total number of interactions, nestedness, and modularity. We found no correlation between any of these network structural variables and PSS values (Fig. **S3**), suggesting that our approach allows for comparisons across different systems with a wide range of network properties. Type I error rates were between 0% and 10% (mean 4%), except for small and equal numbers of rows and columns (< 11 rows × < 11 columns; Fig. **S4a**), small network matrices with unequal numbers of rows and columns (< 15 rows × 20 columns; Fig. **S4b**), and networks with low total number of interactions (< 30 interactions; Fig. **S4c**). This was expected because in these cases most species have a single interaction recorded, which means that their node degree is equal to 1. Consequently, they appear specialized on a phylogenetically clustered lineage.

Rates of Type II error at the CDM level were low for both clustered (1.1%) and overdispersed (5.2%) scenarios. We observed high rates of Type II error (36%) in clustered scenarios when assessed at the plot level, which correspond to single rows in the CDMs.

### Comparison of NRI and PSS

NRI and PSS generally yielded similar results regarding the proportion of taxa and interaction frequencies with random, clustered and overdispersed partners (Table **2**). However, these indices can lead to different results for some datasets. For example, 53.6% of cyanobacterial taxa were found to associate with random partners according to PSS, compared to 17.8% according to NRI (Table **2**). In some cases, such as in the avian seed-dispersal dataset, PSS and NRI inferred a similar percentage of plant species that associate with random partners (38.6% and 37.4%, respectively). However, those species account for different percentages of the interaction frequencies (39.8% and 28.8%, respectively). This indicates that PSS and NRI detected different species that have random partners. These discrepancies are more evident in the comparison of the index values obtained for each taxon (Fig. **3**). These indices yielded highly similar values for some guilds (Fig. **3c, d, f, h**), and very different values for taxa in other guilds (Fig. **3a, b, e, g**). In three cases, the correspondence between values of the two indices was much higher for one of the guilds within the same dataset (e.g., fleas r^2^=0.28 [Fig. **3a**] compared to mammals r^2^=0.74 [Fig. **3b**]). The biggest discrepancies between NRI and PSS were for guilds that are the most dependent on specific partners. This includes obligate parasitic fleas on their mammal hosts, obligate mutualistic fungi on their photoautotrophic and diazotrophic cyanobacteria, and obligate corals on their photoautotrophic dinoflagellate partners (Fig. **3a, e, g**). Even in cases where the two indices inferred the same structure, we observed a slight trend towards more negative values for the NRI index (Fig. **3c, d**). For example, most interaction pairs (dots) from the mammals-fleas dataset fall into the same areas (Fig. **2b, c**) of the NRI-NRI (Fig. **4c**) and the PSS-PSS space (Fig. **4d**). However, the interaction density is shifted towards the left (more negative) according to NRI (Fig. **4c**) compared to PSS (Fig. **4d**). We found a similar result with the avian seed-dispersal dataset, for which NRI inferred a higher density of interactions in the clustered-clustered area (Fig **4g**) compared to PSS (Fig **4h**). In contrast, PSS inferred more negative values than NRI for lichen-forming fungi (Figs **3e**, **4k, l**) and corals (Figs **3g**, **4o, p**).

**Figure 3.**
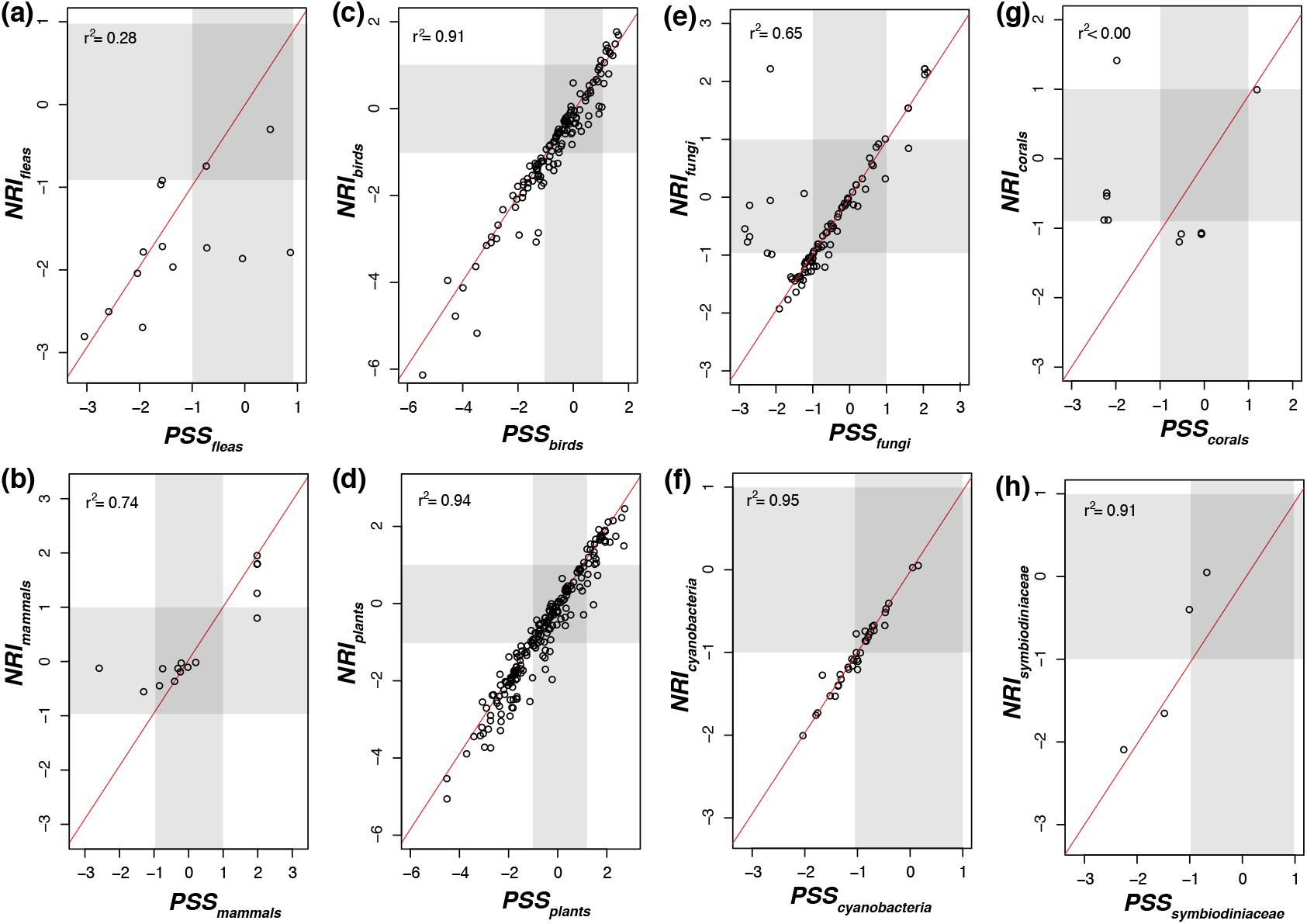
Comparison of NRI and PSS values for the species in each of the eight guilds present in the four empirical datasets we analyzed. If a circle falls along the red diagonal line, it means that the two metrics being compared yield the same value for that particular species. (**a and b**) fleas and their mammalian host, (**c and d**) birds as dispersers of plant seeds, (**e and f**) lichen-forming fungi and their cyanobacterial partners, and (**g and h**) corals and their Symbiodiniaceae partners. The shaded areas represent non-significant clustering or overdispersion. The thresholds of the shaded areas were defined as the mean of the 95% confidence interval of the null distributions generated for each taxon in the datasets. All PSS values were calculated with the availability parameter estimated from interaction frequencies.

**Figure 4.**
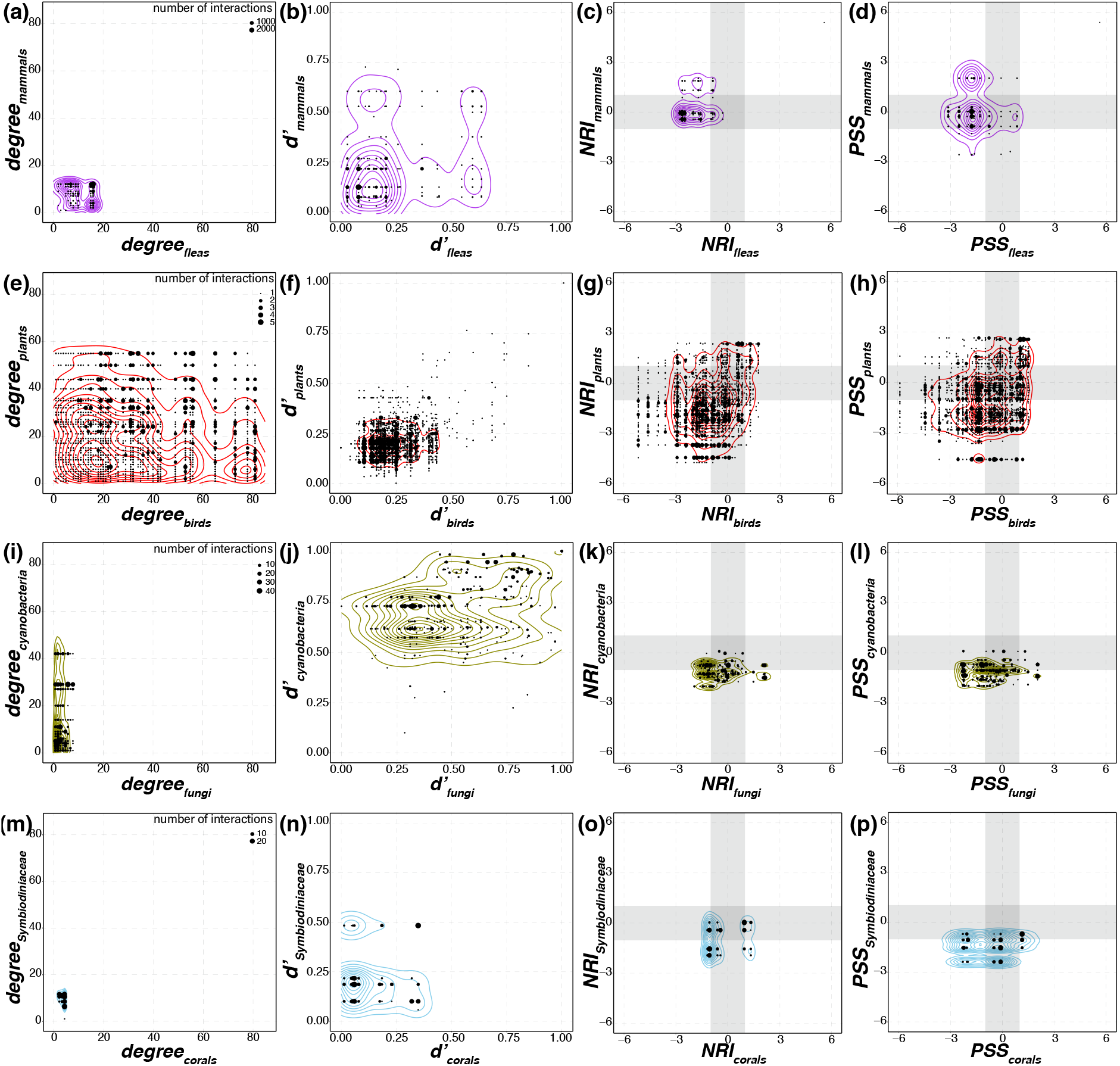
Comparison of biotic co-specialization profiles of four empirical bipartite networks using four metrics (columns): node degree, Blüthgen’s *d’*, NRI and PSS. (**a–d**) Mammals-fleas network from Slovakia (Stanko *et al*., 2002). (**e–h**) Avian seed-dispersal network from the tropical Atlantic forest (Bello *et al*., 2017). (**i–l**) Cyanolichen network from an opportunistic global sampling of the lichen-forming fungal genus *Peltigera* and their *Nostoc* cyanobacterial partners (Chagnon *et al*. 2019). (**m–p**) Corals–dinoflagellates network from the Caribbean (Franklin *et al*., 2012; Fabina *et al*., 2012). Each dot on these plots represents a pair of interacting species. These dots are placed on graphs according to the biotic specificity metric value of the interacting species. For example, in panel **i**, fungi are interacting with one to eight phylogroups of cyanobacteria (X axis) while cyanobacteria are interacting with one to more than 40 fungal species (Y axis) in this network of cyanolichens. All PSS values were calculated with the availability parameter estimated from interaction frequencies. The shaded areas in the panels of the third and fourth columns represent non-significant clustering or overdispersion. The thresholds were defined as the mean of the 95% confidence interval of the null distributions generated for each taxon in the datasets. The size of the dots represents the number of times an interaction was recorded in the matrix, i.e., interaction frequency. Contour lines are estimated 2D distributions.

### Most taxa interact with phylogenetically random or clustered partners

Although we found a wide range of variation in the number of partner species (node degree) in the empirical networks (Fig. **4a, e, i, m**), most interaction pairs involved species without strong specialization signal according to Blüthgen’s *d’* (Fig. **4b, f, n**). The cyanolichen network was an exception, with multiple interaction pairs involving specialist *Nostoc* and generalist (opportunistic) lichenized fungi, or both specialist *Nostoc* and specialist fungi (Fig. **4j**, **2a**). Across all four interaction networks, both NRI and PSS indicated that many taxa interact with random and clustered partners (Table **2**; Fig. **4c, d, g, h, k, l, o, p**). However, taxa that interact with overdispersed partners were rare and not found in all guilds (Table **2**; Fig. **4c, d, g, h, k, l, o, p)**. PSS values for the fleas were highly similar when calculated based on empirical availability or matrix availability (i.e., from interaction frequencies as in equation 4; Fig. **5**).

**Figure 5.**
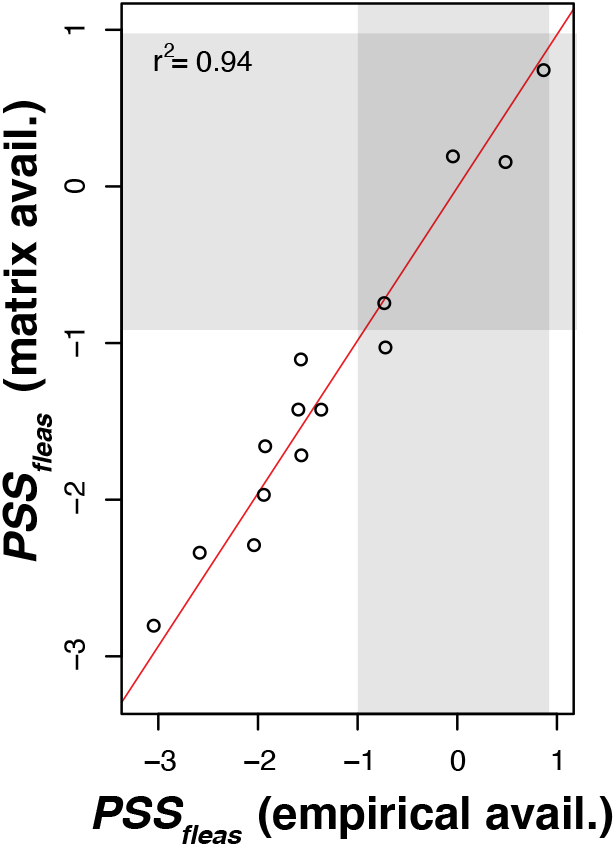
Comparison of PSS values for the mammal partners of flea species estimated using empirical estimates of availability obtained from the environment (X axis), and estimates of availability based on interaction frequencies obtained from an interaction matrix as a proxy for availability from the environment (Y axis). Each circle on these plots represents a species. The thresholds for the shaded areas (random phylogenetic structure) were defined as the mean of the 95% confidence interval of the null distributions generated for each taxon in the datasets. Points on the diagonal red line indicate identical PSS values.

## Discussion

### Advantages of PSS over other indices

PSS integrates both partner availability and phylogenetic structure to characterize biotic specialization of species within interaction networks. As expected when availability is roughly equal among partners (Fig. **1a, d, g**), PSS captures similar information in empirical networks as NRI (Fig. **3c, d, f, h**), an index that accounts for phylogenetic structure without considering partner availability. Therefore, cases where the two indices diverge (Fig. **3a, b, e, g**) are likely due to unequal partner availabilities (e.g., Fig. **1b, c**).

These are situations where existing approaches that do not account for partner availability, such as NRI, can falsely infer clustered or overdispersed phylogenetic structure when the phylogenetic pattern is actually random (i.e., Type I error), or may fail to detect clustering and overdispersion (i.e., Type II error; Kembel, 2009; Miller *et al*., 2017).

The rates of Type I and Type II error observed for PSS are comparable to the best performing combination of phylogenetic diversity metrics + null model as reported in a previous study (regional null + MPD; Miller *et al*., 2017). However, that combination is designed to describe communities with multiple species, which limits its application for interactions networks where many species often interact with few partners (Appendix **S1**). Furthermore, PSS values are not biased by the total number of interactions in the matrix (i.e., *m* in equation 3; Fig. **S3c**), which is the case for an existing specialization index that integrates availability and phylogenetic structure (Jorge *et al*., 2017; Appendix **S1**).

The higher rates of Type II error that we observed at the plot level of the CDMs were also reported by Miller *et al*. (2017) for other indices. The simulation strategy that we implemented to test Type II error is expected to generate the clustered and overdispersed patterns at the scale of the entire simulated arena. Our CDMs are intended to be a representative sample of that arena. Therefore, calculating PSS at the plot level (i.e., single rows of the matrix) is equivalent to taking a much smaller sample of that arena, which explains why the power of the index decreases. Therefore, we expect that the power of PSS will also decrease when interaction networks are under-sampled.

### Phylogenetic structure in empirical networks

The integration of phylogenetic data with interaction networks can provide insights about the relative importance of ecological and evolutionary processes that shape biological communities (Segar *et al*., 2020). Previous studies have shown that many ecological interactions, as well as interaction-related traits, display phylogenetic structure, where closely related species tend to have overlapping sets of partners (Rezende *et al*., 2007; Gómez *et al*., 2010; Eklöf *et al*., 2012; Aizen *et al*., 2016). Based on those findings, it should be common for species to be specialized on phylogenetically clustered partners. However, PSS analyses of four empirical networks showed that many species interact with phylogenetically random partners (Table **2**; Fig. **4d, h, l, p**). Our results suggests that while interaction traits can be conserved across some phylogenetic scales, the assemblage of communities of interacting species at regional and local scales can be constrained by the relative effect of processes other than the evolutionary history of the species (Mello *et al*., 2019; Segar *et al*., 2020), such as the availability of potential partners.

Nevertheless, we also encountered many cases of phylogenetic specialization in all four empirical datasets (Table **2**; Fig. **4d, h, l, p**). For example, in cyanolichens, the peak of the distribution of interactions was found to be in the random-clustered and clustered-clustered regions of the PSS-PSS space (Figs **2c**, **4l**).

These results are consistent with past assessments that *Peltigera* species are most often specialized on generalist, but also on specialist, *Nostoc* phylogroups (Magain *et al*., 2017a). Similarly, Krasnov *et al*. (2012) reported that the fleas in the mammals–fleas dataset showed phylogenetic signal in their host range, which is consistent with our observed distribution of fleas infecting a clustered set of mammal hosts at a regional scale (Figs **2c**, **4d**). However, we also detected mammal species that are infected by phylogenetically overdispersed fleas (Figs **2c**, **4d**). In contrast, the tropical avian seed-dispersal network consists mostly of interactions involving generalist species (Figs 2a, **4f**) that may not require specialized traits, or may be specialized on traits that are not phylogenetically conserved (Bolmgren & Eriksson, 2005; Bello *et al*., 2017; Emer *et al*., 2019). This dataset includes a large proportion (75%) of interactions involving species that associate with phylogenetically random partners (Fig. **4h**). However, the seed-dispersal network also includes the largest proportion (22%) and most striking examples of interactions between species with clustered partners (Figs. **2c**, **4h**).

### Is overdispersion a signature of specialists or generalists?

Studies that have used phylogenetic diversity metrics to characterize biotic specialization often focused on cases where partners were significantly more closely related than expected by chance (but see Maherali & Klironomos, 2007), and considered overdispersion as a signature of generalists (Poulin *et al*., 2011; Cooper *et al*., 2012; Jorge *et al*., 2014). This is because overdispersion indicates that a species associates with distantly related partners. However, in a framework where partner availability is accounted for, a significant phylogenetic structure can only be detected when interaction frequencies are non-random. With PSS, overdispersion means that a species interacts with its partners more than expected by chance, and those partners are more distantly related than expected by chance. This is consistent with high intensity of partner use within a narrow span of a species’ biotic niche and, therefore, should be interpreted as a signature of specialists (Fig. **2c**).

### Availability based on interaction frequencies as proxies for availability of partners in nature

Interaction frequencies in network matrices are commonly used as proxies for partner availability in nature, as evidenced by the widespread use of Blüthgen’s *d’* to quantify specialization (Fründ *et al*., 2016; Arceo-Gómez *et al*., 2020; Buitrón-Jurado & Sanz, 2020). However, this proxy might be inaccurate if the interactions are not sampled systematically, when facultative partners are involved, or when interaction frequencies are independent from the availability of partners in nature. We only had direct empirical estimates of partner availability for the calculation of PSS_fleas_ (Table 1; Stanko *et al*., 2002). In that case, we found high correspondence among PSS values calculated based on empirical and matrix availability (Fig. **5**). The matrix availability proxy might be especially problematic for the *Peltigera–Nostoc* dataset, which was sampled at a global scale in a non-systematic way. In this case, the interaction frequencies may lead to highly inaccurate estimates of the partner availabilities, particularly since *Nostoc* symbionts can be free-living (Nelson *et al*., 2020).

**Table 1.** Summary of features of the four empirical datasets used in this study. Int. freq.: availability is estimated from interaction frequencies.

| Dataset<br>Guild | Total number<br>of interactions | Taxonomic<br>scale | No. of<br>taxa | Phylogenetic distances | Availability | Phylogeny source |
| --- | --- | --- | --- | --- | --- | --- |
| **Mammals-fleas** | 11509 |  |  |  |  |  |
| Mammals |  | Species | 19 | Substitutions per site | Int. freq. and empirical | Bininda-Emonds *et al.* (2007) |
| Fleas |  | Genus | 14 | Substitutions per site | Int. freq. | Zhu *et al.* (2015) |
| **Avian seed-dispersal** | 2474* |  |  |  |  |  |
| Birds |  | Species | 183 | Divergence times | Int. freq. | Emer *et al.* (2019) |
| Plants |  | Species | 270 | Divergence times | Int. freq. | Emer *et al.* (2019) |
| **Cyanolichens** | 1026 |  |  |  |  |  |
| Fungi |  | Species | 155 | Substitutions per site | Int. freq. | Chagnon *et al.* (2019) |
| Cyanobacteria |  | Phylogroups and haplotypes<sup>†</sup> | 95 | Substitutions per site<sup>‡</sup> | Int. freq. | Chagnon *et al.* (2019) |
| **Corals-dinoflagellates** | 334 |  |  |  |  |  |
| Corals |  | Species | 12 | Substitutions per site | Int. freq. | Kitahara *et al.* (2010) |
| Symbiodiniaceae |  | Genus | 5 | Substitutions per site | Int. freq. | LaJeunesse *et al.* (2018) |

\*We limited our sampling to avian-seed dispersal interactions that were recorded as part of network studies to limit sampling biases towards the plants or the birds present in the communities.
<sup>†</sup>Both phylogroups and haplotypes were used as proxies for species (Magain *et al.*, 2017a).
<sup>‡</sup>Sequence pairwise distances, corrected with the General Time Reversible model because we lacked well-resolved and well-supported phylogenies for *Nostoc* and, therefore, could not infer phylogenetic distances directly from branch lengths

### Importance of phylogenetic and spatial scales for interpreting PSS values

Interpretations of PSS values must consider the phylogenetic and spatial scales of the datasets. For example, we found that a large proportion (53.6%) of cyanobacterial taxa associate with random partners (Table **2**). However, this network only includes the interactions with species from a single genus of lichen-forming fungi (*Peltigera*). If we had done the same analysis in the context of all lichen-forming fungi (which span multiple classes of Fungi), the partners of many cyanobacterial taxa would be highly clustered and some would be overdispersed. Likewise, the avian seed–dispersal dataset consists of interactions that were sampled in a single region, the Atlantic Forest of Brazil (Bello *et al*., 2017). Using PSS, we found that 38.6% of the interactions in this dataset involve plants whose seeds are dispersed by phylogenetically random birds (Table 2, Fig. **4h**). These sets of bird seed dispersers are phylogenetically random relative to the pool of species in the Atlantic Forest, but they likely represent a non-random subset of the phylogenetic diversity of these species at larger spatial scales, as shown by a continental-scale study in South America (Mello *et al*., 2019).

**Table 2.**
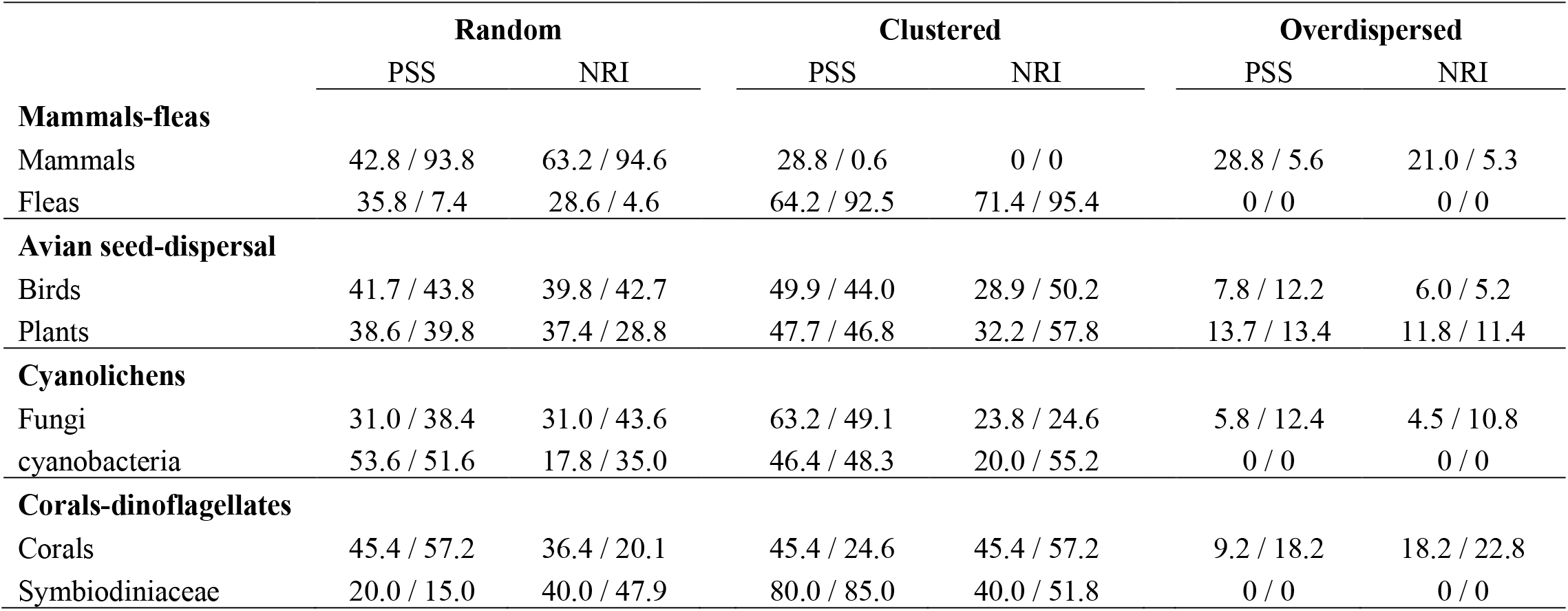
Comparison of PSS and NRI values estimated for taxa across the four empirical datasets used in this study. Note that NRI cannot be calculated for taxa that associate with a single partner. Therefore, we were not able to calculate NRI for 100% of taxa in some datasets. Values before backslash are percentage of taxa, and values after backslash are percentage of interaction frequencies.

### A conceptual framework for an eco-evolutionary interpretation of PSS values

Patterns of phylogenetic diversity are not direct proxies for community assembly processes (Cahill *et al*., 2008; Mayfield & Levine, 2010; Gerhold *et al*., 2015). Instead, we propose testable hypotheses of eco-evolutionary processes that may produce PSS patterns in interaction networks.

Opportunistic interactions can result from multiple processes. Recent colonization or introduction (e.g., long-distance dispersal events or invasive species) into new areas might make opportunistic interactions advantageous in ecological and evolutionary timescales (Poisot *et al*. 2011; Magain *et al*. 2017a). During rapid diversifications, incomplete sorting of traits can generate local populations with high intraspecific variation in interaction traits that allow associations with a broader range of partners. Species may also have spatially structured populations with low phenotypic variation at local scales, but higher variation at larger scales (Batstone *et al*., 2018). This highlights the importance of studying these patterns at multiple spatial scales (Gomulkiewicz *et al*., 2000; Jorge *et al*., 2014). Low heterogeneity in resources exchanged by partners can result in opportunistic interactions (Pinheiro *et al*., 2019). A recent study also showed that high ecological uncertainty can favor generalized host ranges in avian brood parasites (Antonson *et al*., 2020). How and when selection maintains the variation necessary for opportunistic interactions is not fully understood (Vamosi *et al*., 2014; but see Batstone *et al*., 2018), but it seems to be pervasive even in highly intimate symbioses such as lichens and corals (Fig. **4l, p**; Guimarães *et al*., 2007).

Clustered patterns of biotic specificity may arise when the diversification dynamics of one set of organisms is dependent on its interacting partners. In rare cases, this may lead to cospeciation (de Vienne *et al*., 2013). More commonly, clustering results from repeated switches to closely related partners through time (de Vienne *et al*., 2013; Chagnon *et al*., 2019; Thines, 2019) or from the acquisition of a novel partner that promotes speciation of the interacting species, where emerging new species all retain compatibility with the novel partner (Gomulkiewicz *et al*., 2000; Chagnon *et al*., 2019).

Overdispersed patterns of phylogenetic specificity may arise through retention of plesiomorphic traits, convergent evolution, or competitive exclusion of related partners. Coevolutionary theory predicts that convergent evolution of interaction traits is common in mutualistic networks due to indirect selection pressures that spread throughout the networks (Guimarães *et al*., 2011, 2017). However, convergent evolution in interaction networks can also result in random phylogenetic structure if partner compatibility does not systematically evolve on closely or distantly related lineages.

Our approach presents a quantitative and conceptual framework to study specialization in ecological networks and the eco-evolutionary processes that shape it. Furthermore, our PSS index can be used to elucidate the relationship between phylogenetic specialization and the distribution, abundance, and fitness of species in natural communities (Pinheiro *et al*., 2016, 2019). This may have important implications for managing biodiversity when considering species interactions (Harvey *et al*., 2017).

## Acknowledgements

We would like to thank Carine Emer for kindly sharing the seed dispersal datasets with us, and César Quintana-Cataño for discussions that inspired the study. We thank Dominique Gravel and three anonymous reviewers whose critical feedback led to substantial improvements to our manuscript. This study was funded by the National Science Foundation (NSF) awards DEB SG 1556995 and BEE 1929994 to FL and JM, and was supported in part by an NSF Graduate Research Fellowship Program award to IDM under grant no. DGE 1644868. C.J.P-D was also supported by a Special Topics Award from the Mycological Society of America and JPG was supported by the U.S. Department of Energy, Office of Biological and Environmental Research, Genomic Science Program under Award Number DE-SC002036.

## Author Contributions

C.J.P-D. conceived the idea, designed the methodology, developed the code, performed analyses and wrote the manuscript; I.D.M. conceived the idea, designed the methodology, performed analyses and reviewed the manuscript; J.P.G, P.-L.C., N.M. and J.M. provided expertise and feedback, and reviewed the manuscript; F.L. conceived the idea, designed the methodology, provided expertise and feedback and contributed to the writing of the manuscript.

## Data Accessibility Statement

The PSS R package is available at https://github.com/cjpardodelahoz/pss. All datasets, including empirical and simulated interaction matrices and trees, as well as all the code used in this study are available from the Dryad Digital Repository: https://doi.org/10.5061/dryad.s1rn8pk4q.

The following Supporting Information is available for this article:

**Appendix S1** Bias associated with existing methods that incorporate partner availability.

**Appendix S2** Behavior of the index when excluding KL factors ≤ 0.

**Appendix S3** Simulations to asses Type II error.

**Fig. S1** Behavior of two common metrics of phylogenetic diversity across a range of possible number of partner species from a pool of 50 taxa.

**Fig. S2** Variation of network binary connectance with increasing total number of interactions (*m*).

**Fig. S3** Distribution of PSS_*rows*_ on simulated random networks and phylogenetic branch lengths drawn from a lognormal distribution.

**Fig. S4** Distribution of Type I error rates on simulated random networks.

