## Supporting Information for "A new approach to integrate phylogenetic structure and partner availability to study biotic specialization in ecological networks"

Article title: A new approach to integrate phylogenetic structure and partner availability in studies of specialization in ecological networks

Authors: Carlos J. Pardo-De la Hoz, Ian D. Medeiros, Jean P. Gibert, Pierre-Luc Chagnon, Nicolas Magain, Jolanta Miadlikowska, François Lutzoni

The following Supporting Information is available for this article:

**Appendix S1** Bias associated with existing methods that incorporate partner availability.

**Appendix S2** Behavior of the PSS index when excluding KL factors $\leq$ 0.

**Appendix S3** Simulations to assess Type II error.

**Fig. S1** Behavior of two common metrics of phylogenetic diversity across a range of possible number of partner species from a pool of 50 taxa.

**Fig. S2** Variation of network binary connectance with increasing total number of interactions (*m*).

**Fig. S3** Distribution of $PSS_{rows}$ on simulated random networks and phylogenetic branch lengths drawn from a lognormal distribution.

**Fig. S4** Distribution of Type I error rates on simulated random networks.

### Appendix S1 Bias associated with existing methods that incorporate partner availability.

Jorge *et al.* (2014) proposed an approach to incorporate phylogenetic relatedness and partner availability to measure specialization using a phylogenetic diversity metric and Standardized Effect Sizes (SES). A subsequent study by Jorge *et al.* (2017) detected a bias behavior in this approach, as the index values were found to be strongly correlated with interaction frequencies (Fig. 1 in Jorge *et al.*, 2017). We demonstrate here that SES are comparable among species with different interaction frequencies. The bias observed in Jorge *et al.* (2017) arises because the distribution of values that the phylogenetic diversity metrics can take changes as the total number of partners approaches the total number of species included in the phylogenetic tree (Fig. **S1**).

To account for the availability of partners, Jorge *et al.* (2014) proposed to compute the version of the mean pairwise phylogenetic distance (MPD) that is weighted by interaction frequencies (which behaves as in Fig. **S1b**) and compared it to a null distribution where the interaction frequencies are drawn in proportion to the availability of the potential partners (null model 5 in Kembel [2009]). For example, if a focal species is found to have 10 interactions with 2 partners, the null distribution will be computed based on drawing 10 total interactions from the pool of available partners in proportion to their availability. Following this procedure, the null sets will have the same number of interactions as observed in the data, but the number of partners might be different. In fact, unless interactions are fully random, species with a higher number of interactions will be compared against null sets that have a larger set of partners.

As shown in Fig. **S1b**, the values that the weighted MPD can take increase with the number of partner species. Since the index proposed by Jorge *et al.* (2014) is calculated as SES = – ($null_{mean}$ – observed)/$null_{sd}$, the difference between the observed value of the weighted MPD and that of the null model will also become increasingly larger as the number of observed interactions increases. This results in a bias where species that have more interactions deviate more from the null model, when the null model itself cannot take the same values as the observed data. The standard deviation of the null also decreases as the number of partners approaches the total number of taxa in the tree (Fig. **S1b**), which further strengthens the bias.

Jorge *et al*. (2017) document this behavior in their index, which they call Distance-based Specialization Index, or DSI. They proposed a solution to this problem for comparisons among species with different number of interactions. They define DSI*, i.e., a rescaled version of DSI obtained by dividing it by its maximum or minimum possible value within a particular dataset. The computation is shown in equation 2 of Jorge *et al*. (2017):

$$\mathrm{DSI}_i^* = \frac{\mathrm{DSI}_i}{|\mathrm{DSI}_{\mathrm{lim}}|}, \qquad (2)$$

This rescaling bounds DSI* between -1 and 1. The authors argue that those values can now be compared between species. However, this is not the case. The rescaling effectively removes the correlation between the index values and interaction frequencies, but the significance thresholds are different for every value of interaction frequency. Moreover, those thresholds vary non-linearly and are different for each dataset. Therefore, the same DSI* value does not mean the same for different species, because DSI* alone is not suitable for statistical hypothesis testing as it does not inform how much the observed value differs from the null model (Fig. 2 in Jorge *et al*. [2017]). Even if two species have the same number of interactions, the same DSI* value can mean different things if the two species are part of different datasets (Fig. 2 in Jorge *et al*. [2017]).

The authors argue that DSI* values can be interpreted as a continuum between maximum specialization (DSI* = 1) and maximum generalization (DSI* = -1). However, DSI is an SES, not a measurement. Unlike measurements, SES can only be interpreted in relationship to the null model used in their calculation. DSI* alone does not provide information about the null model, and we can only extract it if it is plotted with the significance thresholds. Even then, comparisons between networks are difficult. Furthermore, the maximum and minimum values used to rescale DSI are calculated based on a biased null model, which makes the significance thresholds unreliable as well. Thus, DSI* is unsuitable for comparisons among datasets, and does not provide unequivocal information about the phylogenetic structure of specialization.

In contrast to DSI and DSI*, our index, PSS, does not have a biased distribution dependent on interaction frequencies. We demonstrate this with the results of simulations presented in Fig. **S3c**. The reason why PSS is not biased is because our null model maintains the observed interaction frequencies and the observed number of partners for every focal species. Another alternative to avoid the bias in DSI is to simulate null sets of partners by sampling them in proportion to their availability, and then estimate the null distribution using only the sets that have the same number of partners as observed for a focal species. This way, the SES can be computed using only null expectations that have the same total number of partners as observed for a focal species. This is the essence of the "regional null" proposed by Miller *et al*. (2017) to describe the phylogenetic structure of communities. However, the use of the "regional null" is limited for interaction networks because many species interact consistently with a small subset of the available partners (Olesen *et al*., 2007; Chagnon *et al.,* 2018). In these cases, the probability of sampling sets of interactions with the appropriate number of partners decreases as the number of recorded interactions increases. For species with only one or two partners, the null becomes a trivial randomization of the observed interaction frequencies that is no longer representative of a neutral process that depends only on the availability of the partners (Kembel, 2009).

### Appendix S2 Behavior of the PSS index when excluding KL factors $\leq 0$.

The KL factors are calculated as shown in equation 9 of the main text:

$$\mathrm{KL}_k = \mathrm{P'}_{ik} \ln\left(\frac{\mathrm{P'}_{ik}}{q_k}\right), \qquad (9)$$

where $\mathrm{P'}_{ik}$ is the proportion of interactions of species $i$ with species $k$, relative to the total number of interactions of species $i$. $q_k$ is the proportion of interactions of species k with all species, relative to the total number of interactions in the matrix. In other words, $q_k$ represents the availability of $k$. Therefore, equation 9 is a ratio between the frequency of $i$ interacting with $k$ relative to the overall availability of $k$. Equation 9 takes negative values when a species interacts with a partner in a frequency that is less than expected by chance (i.e., $\mathrm{P'}_{ik}, < q_k$). These situations correspond to random interactions. Therefore, we only consider the partners for which the KL factors are > 0, which correspond to the interaction frequencies that deviate from the expectation (i.e., interactions for which $\mathrm{P'}_{ik} > q_k$). This strategy avoids potential negative values in equation 10 of the main text, which would not be meaningful.

Our strategy is an extension of the way we deal with the partners with which the focal species has 0 interactions. Consider the following hypothetical set of interactions recorded for species S1 and assume that all potential partners (A, B, C, D and E) are equally available, each representing 20% of the pool of available partners:

| | A | B | C | D | E |
|---|---|---|---|---|---|
| S1 | 10 | 0 | 10 | 10 | 10 |

In this situation, 25% of the interactions of species S1 are with A, although A represents only 20% of the partners. This means that S1 is interacting with A slightly more frequently than expected by chance. The same is true for the interactions of S1 with C, D, and E. The interactions between S1 and B represent 0% of the total number of interactions of S1, even though B accounts for 20% of the available partners. The expectation is that 20% of the interactions of S1 would be with B. The KL factor calculated from B is not included in the calculation of the PSS index because B was not recorded as one of the partners. However, this is also a case where the proportion of interactions with a partner is less than expected given its availability. We only want to give weight to interaction frequencies that deviate from the expectation. Therefore, it makes sense to prevent the insertion of negative values by only calculating the KL weights for the interaction pairs that deviate from the expectation, the same way we already do for unseen interactions.

No information is lost by only considering interaction pairs with positive KL weights. This is because the proportions of interaction frequencies are not independent. For example, consider another hypothetical case of a species S2 interacting with the same potential partners A, B, C, D, and E:

| | A | B | C | D | E |
|---|---|---|---|---|---|
| S2 | 10 | 4 | 10 | 10 | 10 |

If we assume again that all partners are equally available (i.e., 20% each), we have a situation where the proportion of interactions of S2 with B is less than expected by chance, but not equal to 0. If we were to calculate KL (equation 9) for the interaction between S2 and B, we would get a negative value. However, even if we do not consider the interactions of S2 with B for the KL weights and the phylogenetic distance, we can still distinguish the specialization of S1 and S2. Although we would be considering interactions that have the same frequency, they do not represent the same proportions. The interactions between S2 and A, C, D, E also deviate from the expectation given the availability, but they deviate less. In the case of S1, each of these interactions represent 25% of the total, and in the case of S2 they represent ~22%. Equation 9 will generate values that are intuitive and reflective of the data for the two species, i.e., that interactions of S1 deviate more from what is expected by chance than those of S2. In other words, S1 is slightly more selective than S2.

### Appendix S3 Simulations to assess Type II error

For all simulations, we first generated a random phylogenetic tree with 50 taxa using the rcoal() function in the R package *ape* (Paradis *et al.*, 2004). To simulate clustered communities, we used the function filteringArena() from the R package *metricTester* (Miller *et al.*, 2017). This simulates spatially explicit communities of individuals drawn from the pool phylogeny by assigning them to a spatial arena (316 × 316 spatial units) depending on two trait values evolved under a Brownian motion model in the phylogeny. The trait values correspond to X and Y coordinates in the arena, which causes related individuals to be clustered in space. Species abundances were drawn from a lognormal distribution with log mean of 7. We then used the function makeCDM() in the R package *metricTester* (Miller *et al.*, 2017), which places random plots within the simulated arena and generates a community data matrix (CDM) from the individuals included in the plots.

To simulate overdispersed communities, we used the function competitionArena() from the R package *metricTester* (Miller *et al.*, 2017). In this case, individuals are first drawn randomly to the arena. Then, a fraction of the individuals is removed such that the mean relatedness within radius of 30 spatial units is maximized. Finally, the same number of individuals, as removed in the previous step, was drawn from the pool into the arena. This process was iterated 100 times for each arena. We then generated a CDM as explained above.

We simulated 1000 CDMs for both clustering and overdispersion, and then calculated PSS on the plots of these CDM. We used Wilcoxon signed rank tests to determine whether the distribution of PSS values for a given CDM was significantly different from 0. We report Type II error rates as the proportion of clustered and overdispersed CDMs where the distribution of PSS values is not significantly smaller or larger than 0, respectively. We also estimated Type II error rates as the proportion of plot-level PSS values for which the observed *klMPD* value was not significantly different ($\alpha = 0.05$) from the null distribution generated as described in equation 11.

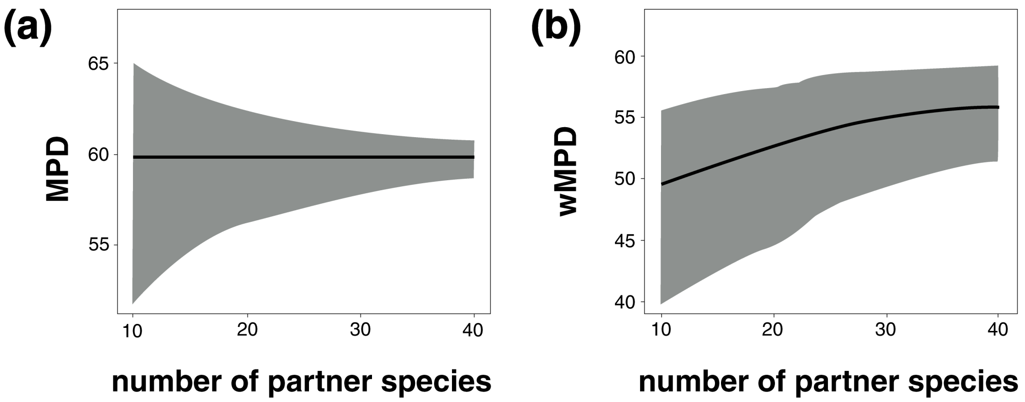


**Fig. S1** Behavior of two common metrics of phylogenetic diversity across a range of possible number of partner species from a pool of 50 taxa. **(a)** MPD: mean pairwise phylogenetic distance of the partners. **(b)** wMPD: MPD weighted by the interaction frequencies. Gray shade shows the distribution of possible values for a given number of partners, and the black line shows the mean of the distribution. Generated as in Miller *et al.* (2017).

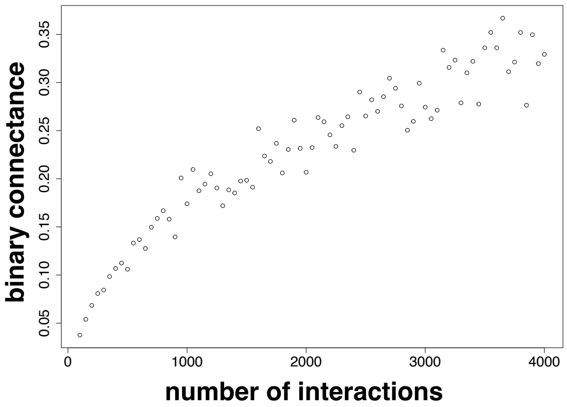


**Fig. S2** Variation of network binary connectance with increasing total number of interactions ($m$). Connectance was calculated for a subset of the 50×50 random matrices simulated for the analysis presented in Fig. **S3c**. We sampled matrices with total number of interactions ranging from 100 to 4000, with increments of 100, for a total 79 matrices.

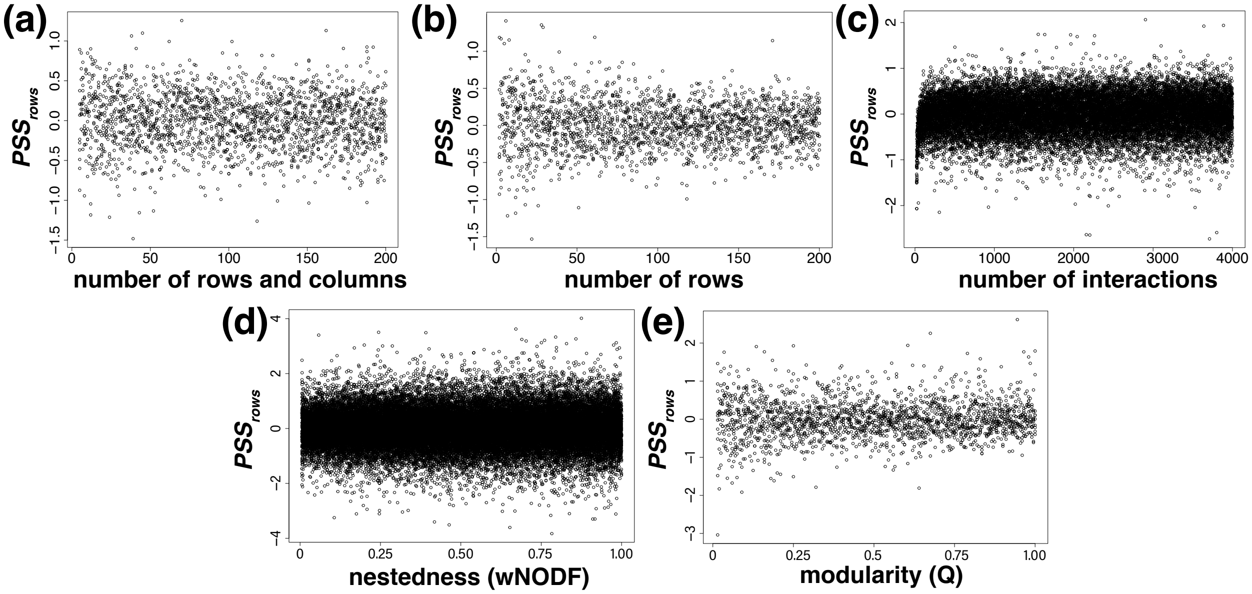


**Fig. S3** Distribution of PSS$_{rows}$ on simulated random networks and phylogenetic branch lengths drawn from a lognormal distribution. Each point on the plots shows the PSS$_{rows}$ value for a simulated matrix on: (a) square random matrices with size varying from 5×5 to 200×200, (b) random matrices with unequal number of rows and columns varying from 2×20 to 200×20, (c) random 50×50 matrices with number of interactions (*m*) varying from 21 to 4000, (d**)** random 50×50 matrices with varying nestedness (wNODF), and (e) random 50×50 matrices with varying modularity (Q).

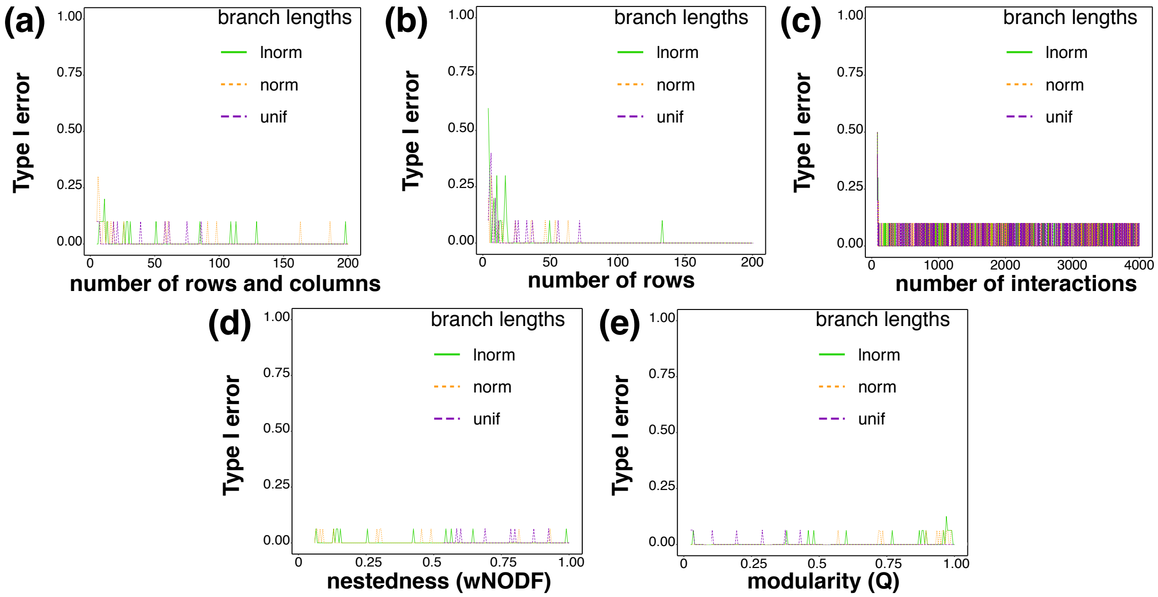


**Fig. S4** Distribution of Type I error rates on simulated random networks with: (**a**) square matrices with sizes varying from 5×5 to 200×200, (**b**) matrices with varying unequal numbers of rows and columns, i.e., varying from 2×20 to 200×20, (**c**) 50×50 matrices with number of interactions (*m*) varying from 21 to 4000, **(d)** 50×50 matrices with varying nestedness (wNODF), and (**e**) 50×50 matrices with varying modularity (Q). lnorm: lognormal, norm: normal, unif: uniform distribution.
